# Niacin Stimulates Mammary Gland Development in Pubertal Mice through Activation of the AKT/mTOR and ERK1/2 Signaling Pathways

**DOI:** 10.1101/2019.12.21.885756

**Authors:** Yu Cao, Juxiong Liu, Lijun Ma, Qing Zhang, Jiaxin Wang, Wenjin Guo, Yanwei Li, Ji Cheng, Shoupeng Fu

**Affiliations:** College of Veterinary Medicine, Jilin University, Changchun 130062, China

**Author notes:** These authors contributed equally to this work (Yu Cao, Juxiong Liu).

**Keywords:** Niacin, mammary gland development, Gi, Akt/mTOR, ERK1/2, pubertal mice

## Abstract

Previous studies have shown the effects of vitamins on the development of the mammary gland. However, the role of niacin in this process has not been reported. Therefore, the aim of this study was to investigate the effects of niacin on mammary gland development in pubertal mice and to use a mouse mammary epithelial cell line to study the underlying mechanism. The results showed that niacin could activate the AKT/mTOR and ERK signaling pathways by the Gi protein-coupled receptor and increase phosphorylation of 4EBP1 to promote the synthesis of cell proliferation markers, leading to the dissociation of the Rb-E2F1 complex in mMECs. In addition, 0.5% niacin promoted mammary duct development, increased the expression of cyclin D1/D3 and PCNA, and activated Akt/mTOR and ERK1/2 in the mammary glands of pubertal mice. These results strongly suggest that niacin stimulates mammary gland development in pubertal mice through the Akt/mTOR and ERK1/2 signaling pathways and that the Gi protein-coupled receptor is essential for this function.

## INTRODUCTION

The mammary gland is the only organ by which mammals feed offspring, and its development determines the health status of the infant. According to the hormonal changes in women, mammary gland development mainly occurs during embryonic development, puberty, pregnancy, lactation and involution[1]. During puberty, periodic estrus causes an increase in ovarian steroid secretion, which in turn stimulates rapid increase in the milk duct branch in mice. Proliferation of mammary gland terminal bud (TEB) cells causes the duct of the mammary gland to attack the outer edge of the fat pad[2]. Moreover, ductal morphogenesis directly determines the number of alveolar cells during lactation[3]. Therefore, ensuring normal development of the mammary gland in adolescent women is the basic condition for maintaining lactation.

The mammary gland undergoes the formation of TEBs and ductal branching in response to estrogen and progesterone produced during the estrous cycle. Estrogen and progesterone induce epithelial cell proliferation mainly by activating the AKT and MAPK signaling pathways[4]. However, in the absence of certain nutrients, such as long-chain fatty acids and amino acids[6-9], the mammary gland does not respond to estrogen stimulation. However, studies on essential vitamins are rare, and some studies have reported that vitamins may have a regulatory effect on mammary gland development; for example, knockout of the vitamin D3 receptor VDR results in the milk ducts overgrowing[10]. Vitamin A acid acts as an intermediate metabolite of vitamin A in animals and has a certain regulatory effect on the moderate growth of the mammary gland[11].

Niacin, an essential vitamin in organisms, plays an important regulatory role in lipid metabolism[12]. It is known that niacin can inhibit lipolysis of triglycerides in adipocytes, thereby inhibiting the release of free fatty acids from adipocytes and reducing the content of free fatty acids in plasma[13]. Niacin can also reduce the synthesis of very low-density lipoprotein and low-density lipoprotein particles in the liver and increase high-density lipoprotein levels[14]. Moreover, niacin displays multiple anti-inflammatory properties in a variety of tissues[15, 16]. Niacin promotes the synthesis of growth hormone[17, 18]. In addition, previous studies have shown that niacin can promote the development of immature follicular cells and the proliferation of cancer cells[19-21].

In summary, niacin has important regulatory effects in both physiological and pathological processes, but the effects of niacin on mouse mammary gland development have not been reported. Therefore, this study aimed to investigate the effect of niacin on the proliferation of the mouse mammary epithelial cell line EPH4EV and the development of the mouse mammary gland. In addition, this study also addressed the underlying mechanisms of this process, including the intrinsic signal transduction pathway.

## MATERIALS AND METHODS

### Chemicals and Antibodies

Niacin (72309, purity ≥99.5%) was purchased from Sigma-Aldrich (USA). DAPI staining solution, the cell counting kit-8 and the cell cycle and apoptosis analysis kit were purchased from Beyotime Biotechnology Inc(China). The Pierce™ class magnetic IP/Co-IP kit was purchased from Thermo Scientific Biotechnology Inc (USA). The 5-ethynyl-2′-deoxyuridine (EdU) incorporation assay kit was purchased from RIBOBIO Biotechnology Co., Inc (China). DMEM (Dulbecco’s modified Eagle’s medium) was purchased from Gibco (USA). FBS (fetal bovine serum) was purchased from Clark (USA). The selective inhibitors MK2206 (AKT), U0126 (ERK1/2), and rapamycin (mTOR) were purchased from Selleck Chemicals (USA).

Polyclonal antibodies against ERK1/2 (16443-1-AP), AKT (10176-2-AP), AKT-phospho-S473 (66444-1-lg) and Cyclin D1 (60186-1-lg) were purchased from Proteintech Co., Inc. (China). Polyclonal antibodies against 4E-BP1 (53H11), p-Rb (Ser807/811) (D20B12), cyclin D3 (DCS22), p-ERK, mTOR, p-mTOR, TUB, and phospho-4E-BP1 (Thr37/46) (236B4) were purchased from Cell Signaling Technology Inc. (USA). Rabbit IgG (A7016) was purchased from Beyotime Biotechnology Inc. (China). Polyclonal antibodies against PCNA (sc-25280) were purchased from Santa Cruz Biotechnology (USA). Polyclonal antibodies against Rb (ab181616) and E2F1 (ab179445) were purchased from Abcam (UK). The related secondary antibodies were purchased from BOSTER Technology, Inc. (USA).

### Cell Culture and Treatment

mMECs (EPH4EV CELL) purchased from American Type Culture Collection (ATCC CRL-3063TM) were cultured in DMEM containing 10% FBS at 37°C in a humidified incubator at 5% CO_2_. Then, 25,000 cells were added to a 60×15 mm dish, and different concentrations of niacin (0, 100 μM, 500 μM, 1 mM, 1.5 mM, and 2 mM) were added to each dish and cultured for 2 d to study the role of niacin in mMECs. The effects of the Gi protein-coupled receptor and AKT, ERK and mTOR signaling pathways on niacin-mediated mMEC proliferation were studied by treating mMECs with 1 mM niacin in the presence or absence of PTX, MK2206, U0126 and rapamycin for 2 d. The culture medium was changed once per day.

### CCK8 and EdU Assays

After the mMECs were incubated with niacin for 2 d, mMEC proliferation was detected using CCK8 and EDU kits according to the manufacturer’s instructions. Briefly, the mMECs were incubated at 37°C with 10 μL of CCK8 in each well for 1 h. Subsequently, absorbance (OD) was measured at 450 nm on a microplate reader (USA). For the EdU incorporation assay, the mMECs were incubated with 50 μM EdU for 0.5 h. After washing 3 times with PBS, the cells were fixed with 4% paraformaldehyde for 12 h, stained with Apollo solution for 30 min in the dark and stained with DAPI for 3 min. The images were observed under a fluorescence microscope (TCS SP5; Leica, Mannheim, Germany).

### Immunoprecipitation assay

Cell supernatant was collected in a 1.5 mL centrifugal tube using lysis buffer (0.025 M Tris, 0.15 M NaCl, 0.001 M EDTA, 1% NP 40 and 5% glycerol), and Rb or pRb antibodies were added to each sample to form immune complexes. Then, 25 μL Pierce protein A/G was added to each sample tube, and the magnetic beads were collected by a 12-tube magnetic separation rack (CST). Finally, 100 μL eluent was added to elute the proteins on the magnetic beads, and the samples were prepared for Western blot experiments.

### Western blot analysis

The total proteins of mMECs and mouse mammary glands were isolated using RIPA lysis buffer, and the protein concentrations were determined using a Pierce™ BCA protein assay kit. Subsequently, the total proteins were separated by SDS-PAGE and transferred to a PVDF membrane. The PVDF membrane was incubated overnight at 4° with primary antibody (1:1,000 dilution), and then the membrane was incubated for 1 h at room temperature using HRP-conjugated goat anti-mouse (1:3,000) or goat anti-rabbit secondary antibody (1:3,000). The protein bands were visualized using an ECL Plus hypersensitive luminescent solution according to the manufacturer’s instructions.

### Real-time quantitative PCR

After the mMECs were treated with niacin for 2 d, the mRNA expression levels of CDC6, ESPL1, and POLE2 were detected by real-time quantitative PCR. Total RNA was extracted from mMECs using TRIzol reagent in accordance with the manufacturer’s instructions. Two micrograms of RNA was reverse transcribed into cDNA. Real-time quantitative PCR was performed on a CFX96 system using SYBR Green Supermix. β-actin was used as a reference gene, and the amount of mRNA was calculated with the 2^-ΔΔCT^ method. All real-time quantitative PCR analyses were repeated 3 times. The primer sequences were as follows as Table 1.

**Table 1.**
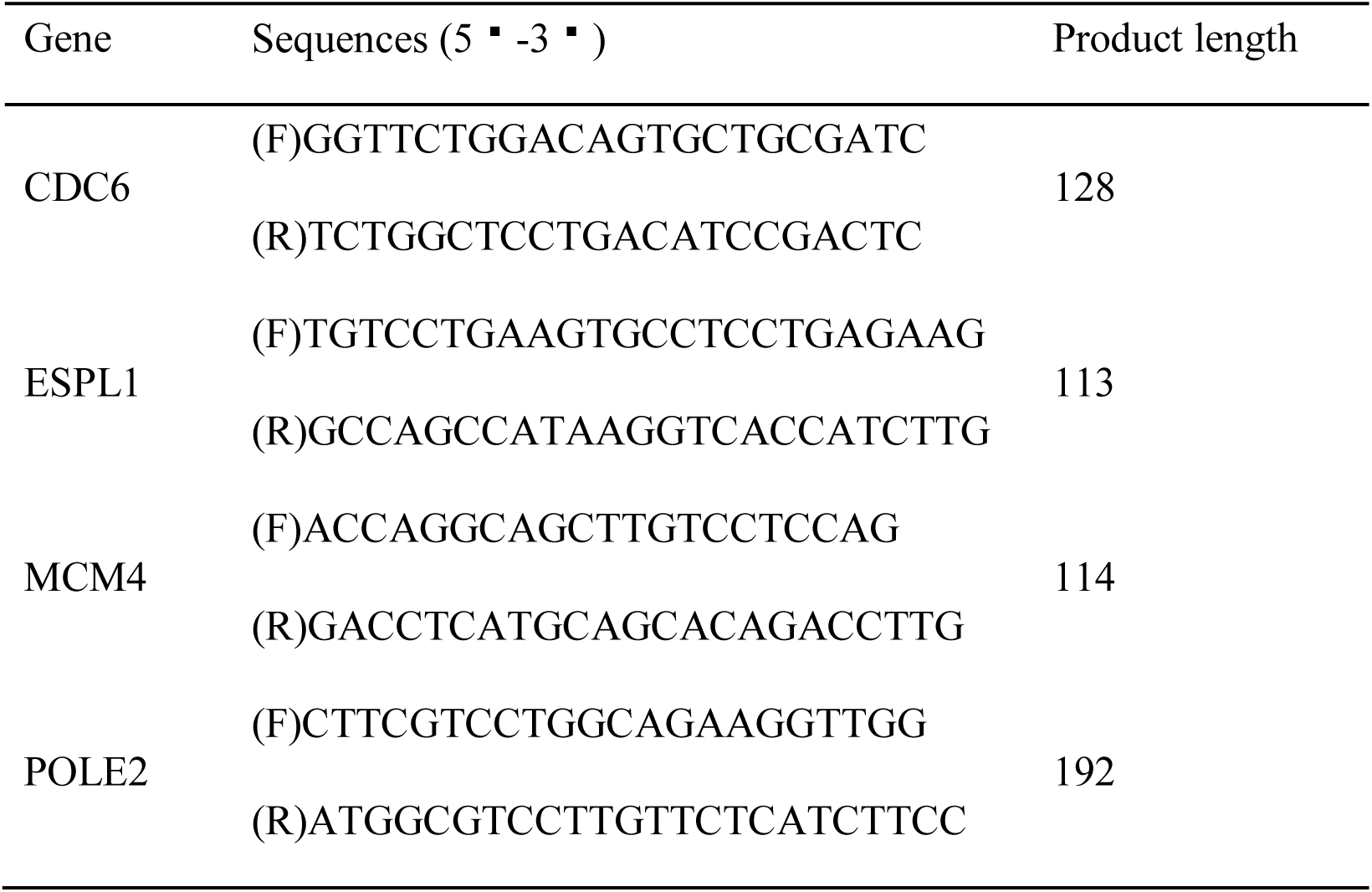
Primer sequences of CDC6, ESPL1, MCM4 and POLE2.

### Flow cytometry analysis

The mMECs were trypsinized and transferred to a 1.5 mL centrifuge tube, fixed with 1 mL of precooled 70% alcohol at 4□ for 12 h, and centrifuged at 1,000 g for 3-5 min. The supernatant was carefully removed, retaining approximately 50 μL 70% ethanol. One milliliter of precooled PBS was added, and the cells were resuspended. The cells were centrifuged, the supernatant was carefully removed, and 50 μL of PBS was retained. The bottom of the tube was flicked to disperse the cells to avoid cell aggregation. Then, 0.5 mL propidium iodide staining solution was added to each sample, and the cells were slowly and fully resuspended and kept at 37°C for 30 min in the dark. Cell cycle was detected using a flow cytometer (BD LSRFortessa™).

### Whole mount staining

The mice were sacrificed by cervical dislocation. After the limbs were fixed, the skin was opened, and the peritoneum was exposed. The skin was bluntly opened to expose the mammary gland. The mammary gland was removed and plated on 4% polylysine-treated slides and fixed overnight using Carnoy’s fluid. The mammary glands were immersed in 70%, 35%, and 15% ethanol and distilled water for 10 min and stained with carmine alum (2 g/L) at room temperature overnight. The slides were rinsed in alcohol for 10 min after the overnight staining. The slides were then immersed in xylene, and the results were observed using a stereomicroscope.

### Animals and in Vivo Study

All animal experiments and care procedures were carried out in accordance with the Jilin University Institutional Animal Care and Use Committee (approved on 27 February 2015, Protocol No. 2015047). Pregnant mice were housed in an environmentally controlled room with free access to food and water during a 12-hour light-dark cycle. After the mice were born, they were taken care of by the mother for 4 weeks. After 4 weeks, the mice were divided into the control group (fed the control diet) and the niacin group (drinking water contained 0.5% niacin). After 3 weeks, the mice were sacrificed, and the 4th pair of mammary glands was collected. The left mammary gland was frozen at −80°C for further analysis. The right breast was used for tissue staining.

### Statistical analysis

All data are presented as the means ± standard error of the mean (SEM). Differences between means were determined using Student’s t-test and one-way ANOVA, and a confidence level of P < 0.05 was statistically significant.

## RESULTS

### Niacin promotes proliferation of mMECs by regulating the expression of proliferation markers

To study the proliferative effect of niacin on mMECs, cells were cultured in DMEM and supplemented with different concentrations of niacin (0, 100 μM, 500 μM, 1 mM, 1.5 mM, and 2 mM) for 2 d, and 1 mM niacin had the most obvious effect on the proliferation of mMECs (Figure 1A). Therefore, 1 mM niacin was selected and used in subsequent studies. Niacin significantly increased the protein expression of cyclin D1/D3 and PCNA in mMECs (Figure 1B-E). In addition, flow cytometry analysis showed that 1 mM niacin significantly increased the number of cells in the S and G2/M phases and reduced the number of cells in the G1/G0 phase (Figure 1F). These results indicate that niacin promotes the proliferation of mMECs by regulating the expression of proliferation markers.

**FIG 1.**
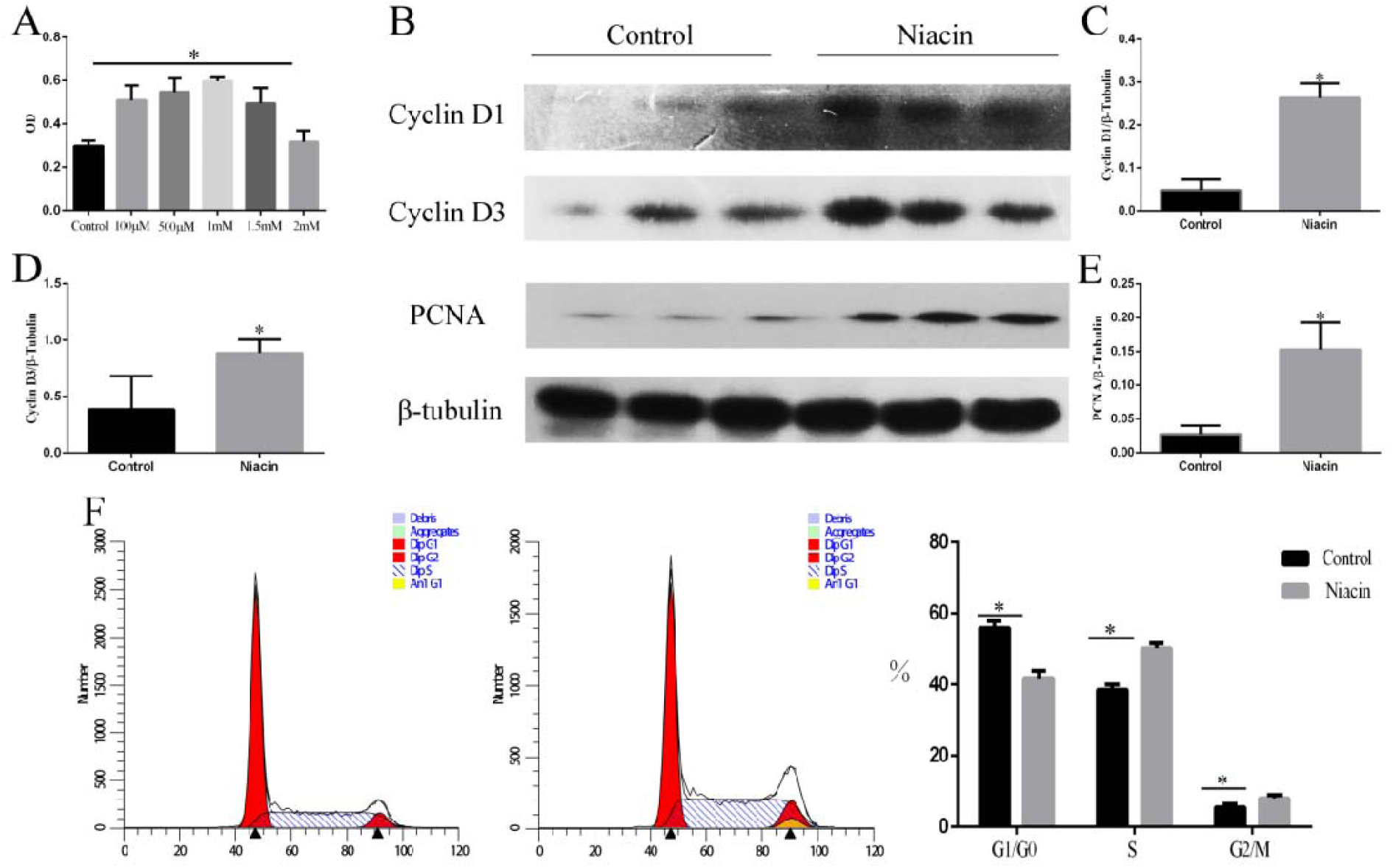
Niacin promotes mMEC proliferation and increases the expression of cell proliferation markers. The mMECs were treated with 1 mM niacin for 2 d. (A) The effect of various concentrations of niacin (0, 0.1, 0.5, 1, and 2 mM) on the proliferation of mMECs was determined by CCK8 analysis. (B) Cyclin D1/D3 and PCNA protein levels were detected by Western blotting. (C, D, E) Immunoblot bands of cyclin D1/D3 and PCNA were digitized and are expressed as the ratios to β-tubulin. Data are expressed as the mean ± SEM. (F) Cell cycle progression was examined by flow cytometry analysis. *P < 0.05 versus the control group.

### Inhibition of Akt/mTOR completely blocks the promotion of mMEC proliferation induced by niacin

To further demonstrate the role of the AKT/mTOR signaling pathway in niacin-induced proliferation of mMECs, the AKT- and mTOR-specific inhibitors MK2206 and rapamycin were used to inhibit the phosphorylation of AKT and mTOR, respectively. The results showed that the significant increases in the p-Akt/Akt and p-mTOR/mTOR ratios in response to 1 mM niacin were reversed by 10 μM MK2206 and 200 nM rapamycin, respectively (Figure 2A, E, and F and Figure 3A and E). Moreover, the EDU and CCK8 results showed that MK2206 and rapamycin abolished the proliferation of mMECs (Figure 2G-I 3F-H). Furthermore, MK2206 and rapamycin reversed the effects of niacin on the protein expression of cyclin D1/D3 and PCNA (Figure 2A-D, Figure 3A-E). These results indicate that niacin promotes the proliferation of mMECs via the AKT/mTOR signaling pathway.

**FIG 2.**
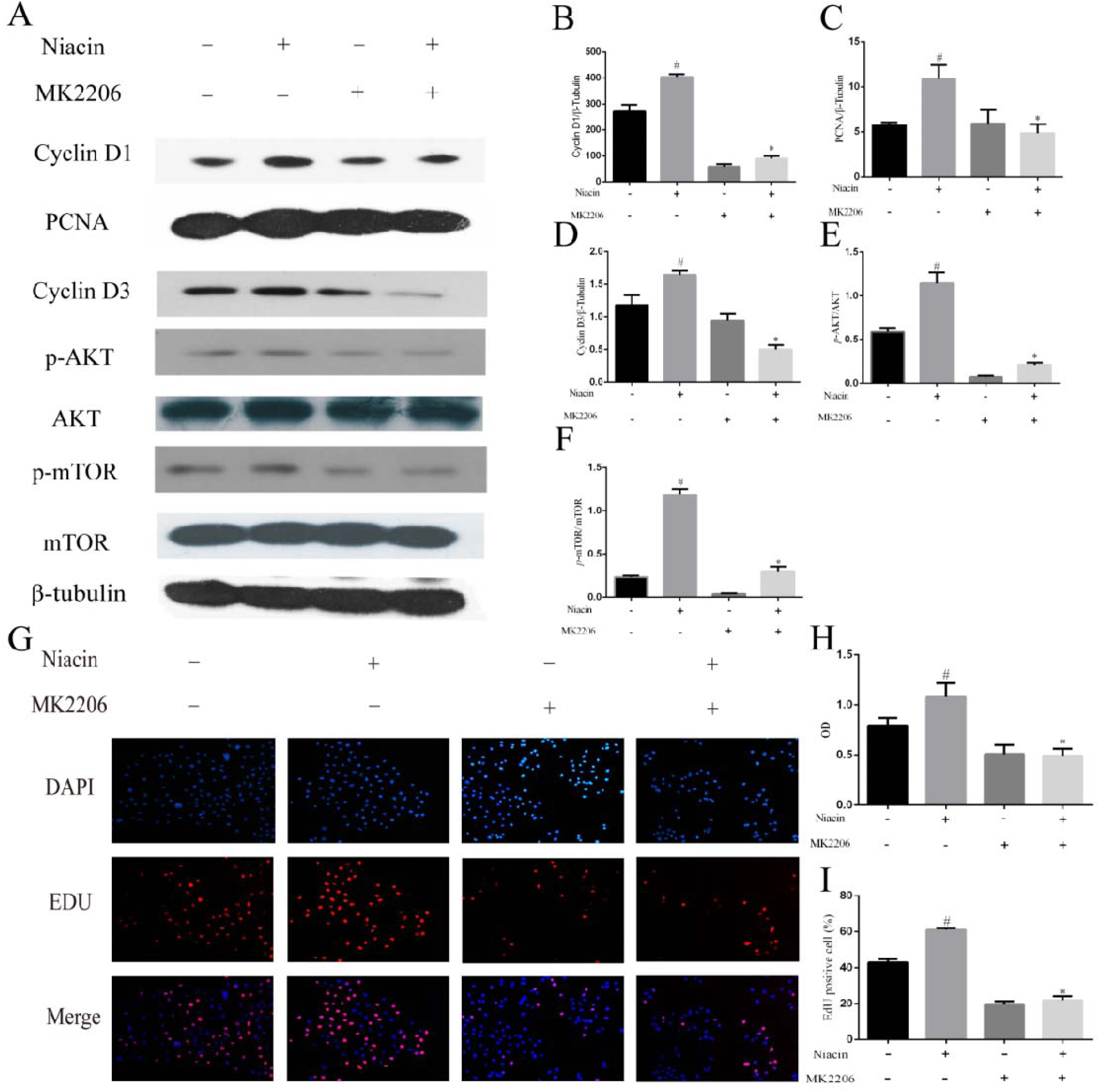
Inhibition of AKT blocks niacin-induced proliferation of mMECs. The mMECs were pretreated for 1 h with MK2206 (2.5 μM) and then incubated for 2 d with niacin (1 mM). (A) The protein levels of p-AKT, AKT, p-mTOR, mTOR, Cyclin D1, PCNA and Cyclin D3 were examined by Western blotting. (B, C, D, E, F). Immunoblot bands of p-AKT/AKT, p-mTOR/mTOR, cyclin D1/D3 and PCNA were digitized, and cyclin D1/D3 and PCNA were expressed as the ratios to β-tubulin. Data are expressed as the mean ± SEM. (G, H, I) The proliferation of mMECs was examined by EdU and CCK8 assays. ^#^P < 0.05 versus the no-treatment group and *P < 0.05 versus the niacin group.

**FIG 3.**
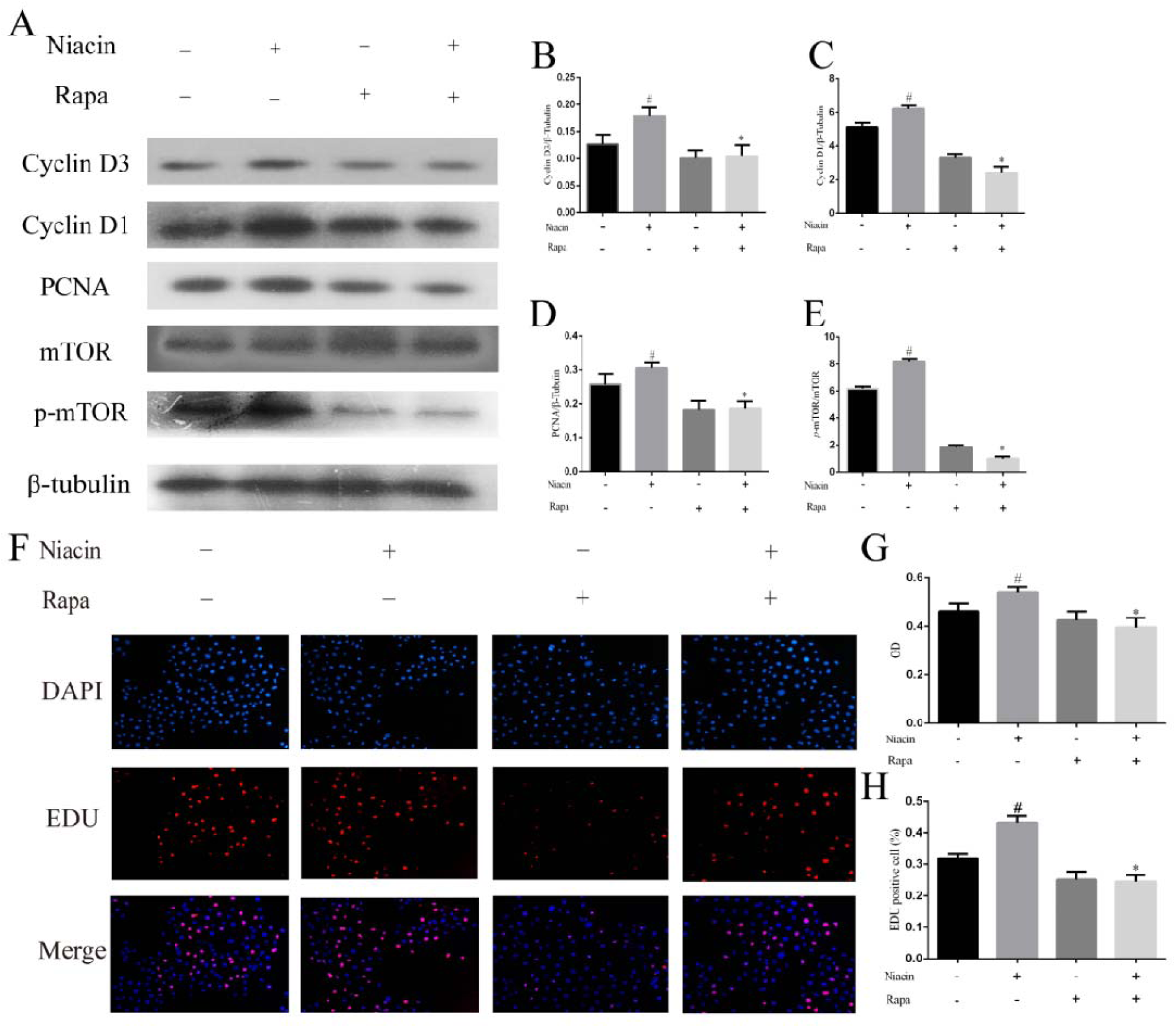
Inhibition of mTOR blocks niacin-induced proliferation of mMECs. The mMECs were pretreated for 1 h with rapamycin (Rapa, 200 nM) and then incubated for 2 d with niacin (1 mM). (A) The protein levels of p-mTOR, mTOR, Cyclin D1, PCNA and Cyclin D3 were examined by Western blotting. (B, C, D, E) Immunoblot bands of p-mTOR/mTOR, Cyclin D1, PCNA and Cyclin D3 were digitized, and Cyclin D1, PCNA and Cyclin D3 were expressed as the ratios to β-tubulin. Data are expressed as the mean ± SEM. (F, G, H) The proliferation of mMECs was examined by EdU and CCK8 assays. ^#^P < 0.05 versus the no-treatment group and *P < 0.05 versus the niacin group.

### Inhibition of ERK1/2 Completely Blocks the Promotion of Niacin-induced mMEC Proliferation

To further demonstrate the role of the ERK1/2 signaling pathway in niacin-induced proliferation of mMECs, the ERK1/2-specific inhibitor U0126 was used to inhibit ERK1/2 phosphorylation. The results showed that the significant increase in the p-ERK/ERK ratio in response to 1 mM niacin was reversed by 10 μM U0126 (Figure 4A, E). Moreover, the results of the EdU and CCK8 assays showed that U0126 blocked the effect of niacin on the proliferation of mMECs (Figure 4F-H). Furthermore, U0126 reversed the effects of niacin on the protein levels of cyclin D1/D3 and PCNA (Figure 4D-G). These results indicate that niacin also promotes the proliferation of mMECs via the ERK1/2 signaling pathway.

**FIG 4.**
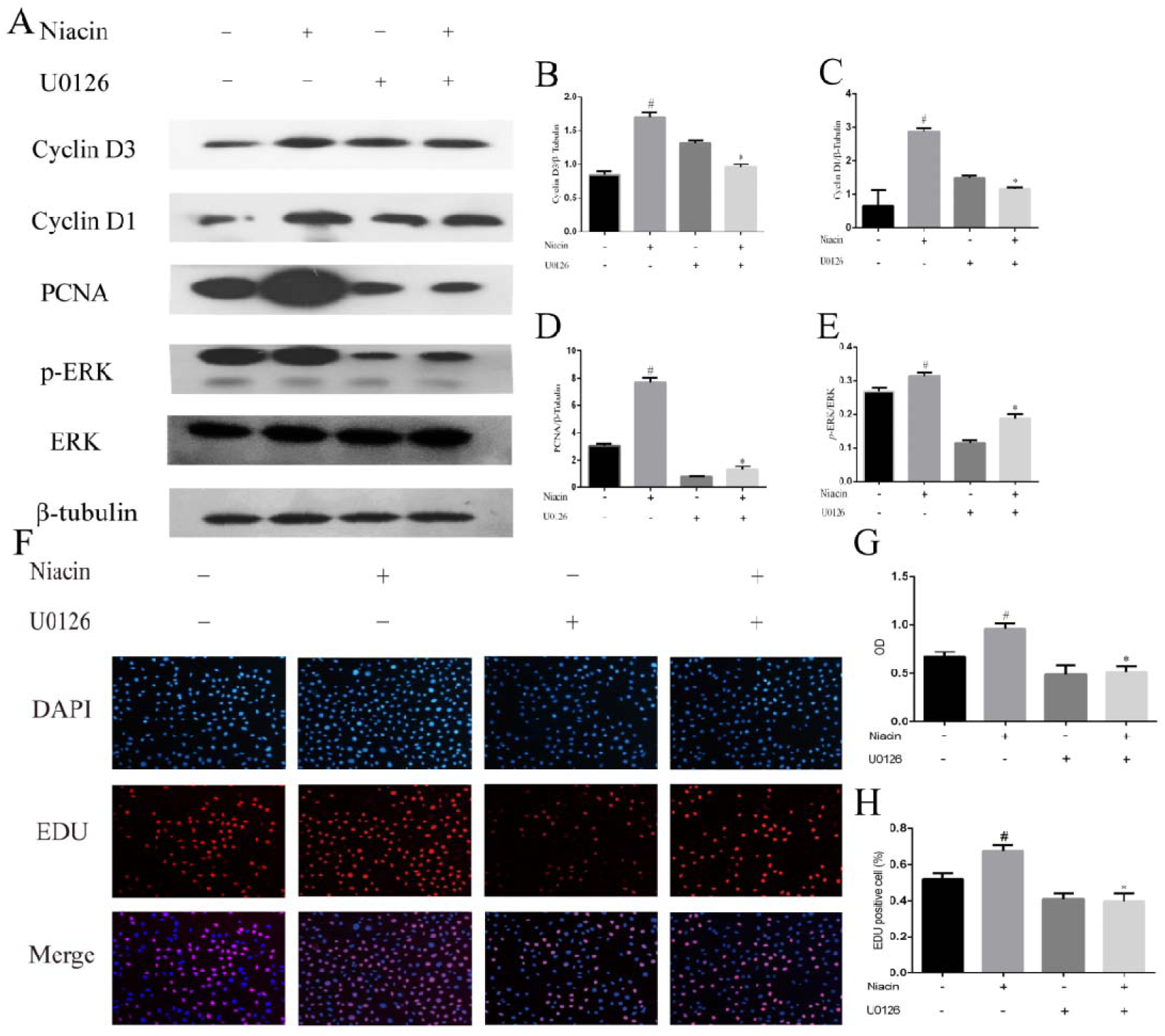
Inhibition of ERK blocks niacin-induced proliferation of mMECs. The mMECs were pretreated for 1 h with U0126 (10 μM) and then incubated for 2 d with niacin (1 mM). (A) The protein levels of p-ERK, ERK, cyclin D1/D3 and PCNA were examined by Western blotting. (B, C, D, E) Immunoblot bands of p-ERK/ERK, Cyclin D1/D3 and PCNA were digitized, and Cyclin D1/D3 and PCNA were expressed as the ratios to β-tubulin. Data are expressed as the mean ± SEM. (F, G, H) The proliferation of mMECs was examined by EdU and CCK8 assays. ^#^P < 0.05 versus the no-treatment group and *P < 0.05 versus the niacin group.

### PTX Eliminates the Enhancement of Niacin-induced mMEC Proliferation

Previous studies have shown that niacin exerts its biological functions through a Gi protein-coupled receptor[16, 21]. To test whether niacin promotes cell proliferation in mMECs via the Gi protein-coupled receptor, the Gi protein inhibitor PTX was used to block the Gi protein-coupled receptor, and cell proliferation and activation of the AKT/mTOR and ERK1/2 signaling pathways were detected. The results of the CCK8 assay showed that after the addition of PTX, the effect of niacin on the proliferation of mMECs disappeared (Figure 5A). Western blot results showed that PTX blocked niacin-induced activation of the AKT/mTOR and ERK1/2 signaling pathways (Figure 4A-E). These results suggest that niacin promotes the proliferation of mMECs through the Gi protein-coupled receptor.

**FIG 5.**
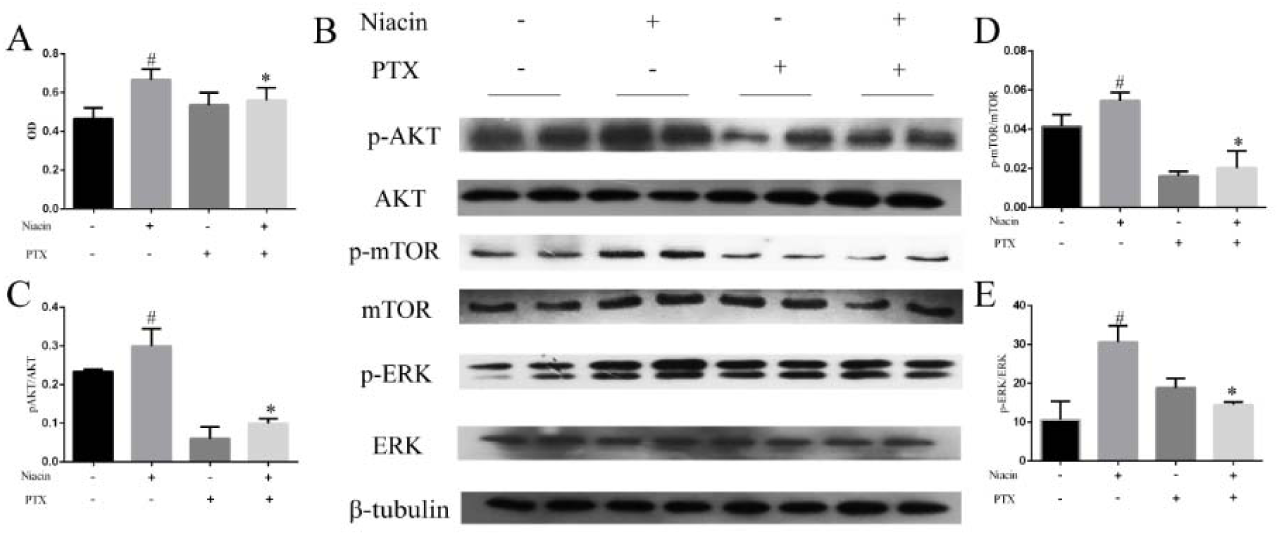
Niacin activates the AKT/mTOR and ERK1/2 signaling pathways via the Gi protein-coupled receptor. The mMECs were pretreated for 1 h with PTX (1 μM) and then incubated for 2 d with niacin (1 mM). (A) The proliferation of mMECs was examined by CCK-8 assay. (B) The protein levels of p-AKT, AKT, p-mTOR, mTOR ERK and p-ERK were examined by Western blotting. (C, D, E) Immunoblot bands of p-AKT/AKT, p-mTOR/ mTOR and p-ERK/ERK were digitized. Data are expressed as the mean ± SEM. ^#^P < 0.05 versus the no-treatment group and *P < 0.05 versus the niacin group.

### Niacin activates 4EBP1 and RB via the Akt/mTOR and ERK1/2 signaling pathways

It has been reported that AKT/mTOR and ERK1/2 signaling pathways can promote downstream 4EBP1 phosphorylation through crosstalk, thereby increasing the expression of cyclin D. Therefore, we used mTOR and ERK1/2 inhibitors (Rapamycin and U0126) to study whether niacin also promotes protein expression through crosstalk of the AKT/mTOR and ERK1/2 signaling pathways. The results showed that niacin significantly increased the ratios of p-4EBP1/4EBP1 and p-Rb/Rb and significantly increased the expression of E2F1 protein. U0126 and rapamycin eliminated the effect of niacin on the increase in p-4EBP1/4EBP1 and p-Rb/Rb ratios (Figure 6A, B C). In addition, U0126 and rapamycin also eliminated the effect of niacin on the expression of E2F1 protein (Figure 6A, D).

**FIG 6.**
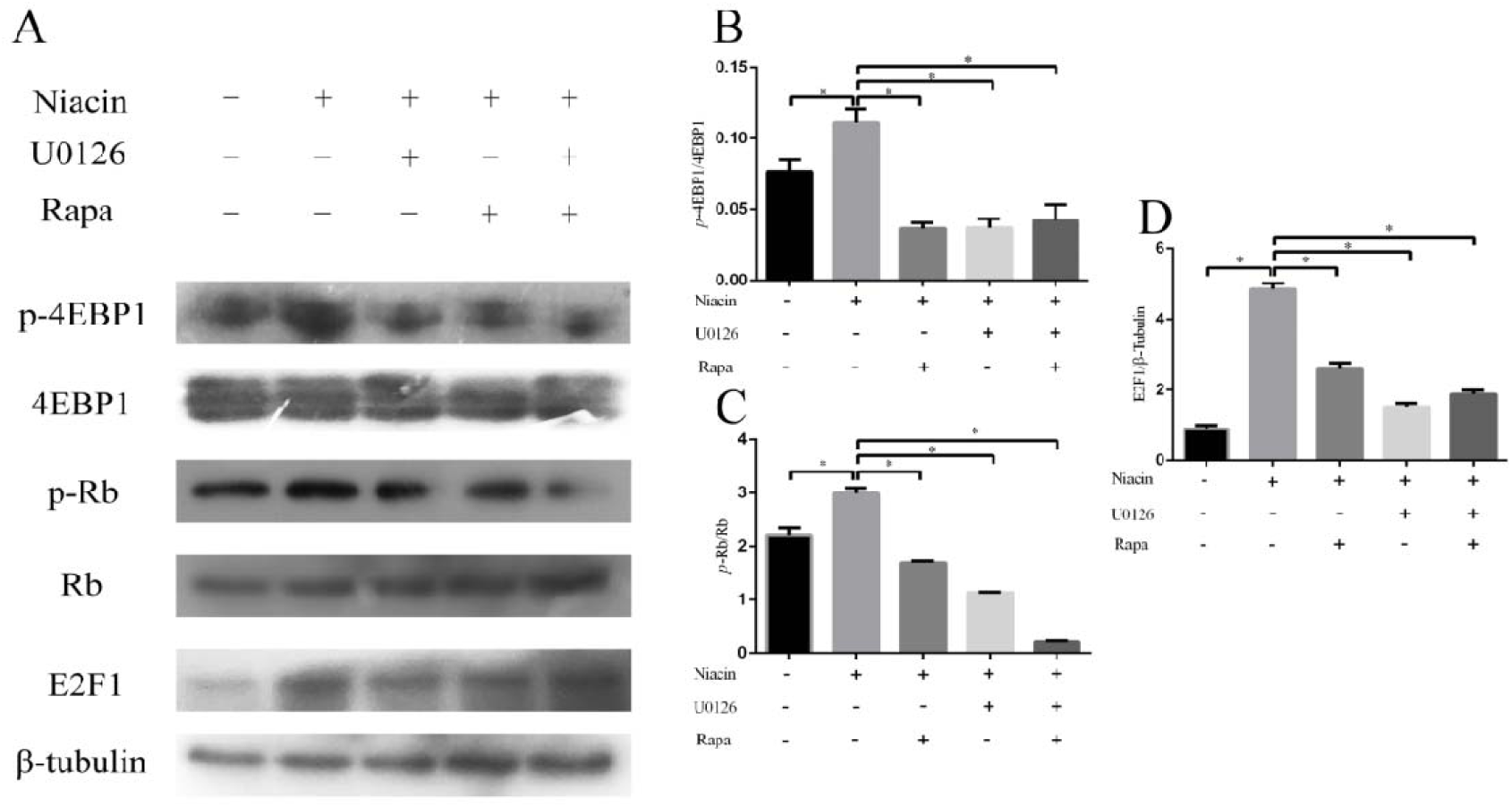
Niacin activates 4EBP1 via the mTOR and ERK signaling pathways. The mMECs were pre-treated for 1 h with rapamycin (200 nM) and U0126 (10 μM) and then incubated for 2 d with niacin (1 mM). (A) The protein expression of p-4EBP1, 4EBP1, p-Rb, Rb and E2F1 was examined by Western blotting. (B, C, D) Mean ± SEM of immunoblot bands of p-4EBP1/4EBP1, p-Rb/Rb and E2F1. *P < 0.05 indicates that the difference is significant.

### Niacin stimulates dissociation of Rb-E2F1 complexes and enhances transcription of E2F1 target genes

Previous studies have shown that cyclin D promotes Rb phosphorylation by binding to cyclin-dependent kinases. Phosphorylation of Rb promotes Rb-E2F1 complex separation and E2F1 target gene transcription, which promotes cell cycle progression and cell proliferation[22, 23]. To test whether niacin increases the release of Rb-bound E2F1 by enhancing the phosphorylation of Rb, an IP assay was carried out. The results showed that the level of the Rb-E2F1 complex decreased significantly after niacin treatment, while the level of the Cyclin D1-Rb complex increased significantly (Figure 7A). These results suggest that after niacin stimulation, Rb phosphorylation is increased by binding to cyclin D and subsequently leads to E2F1-RB complex dissociation.

**FIG 7.**
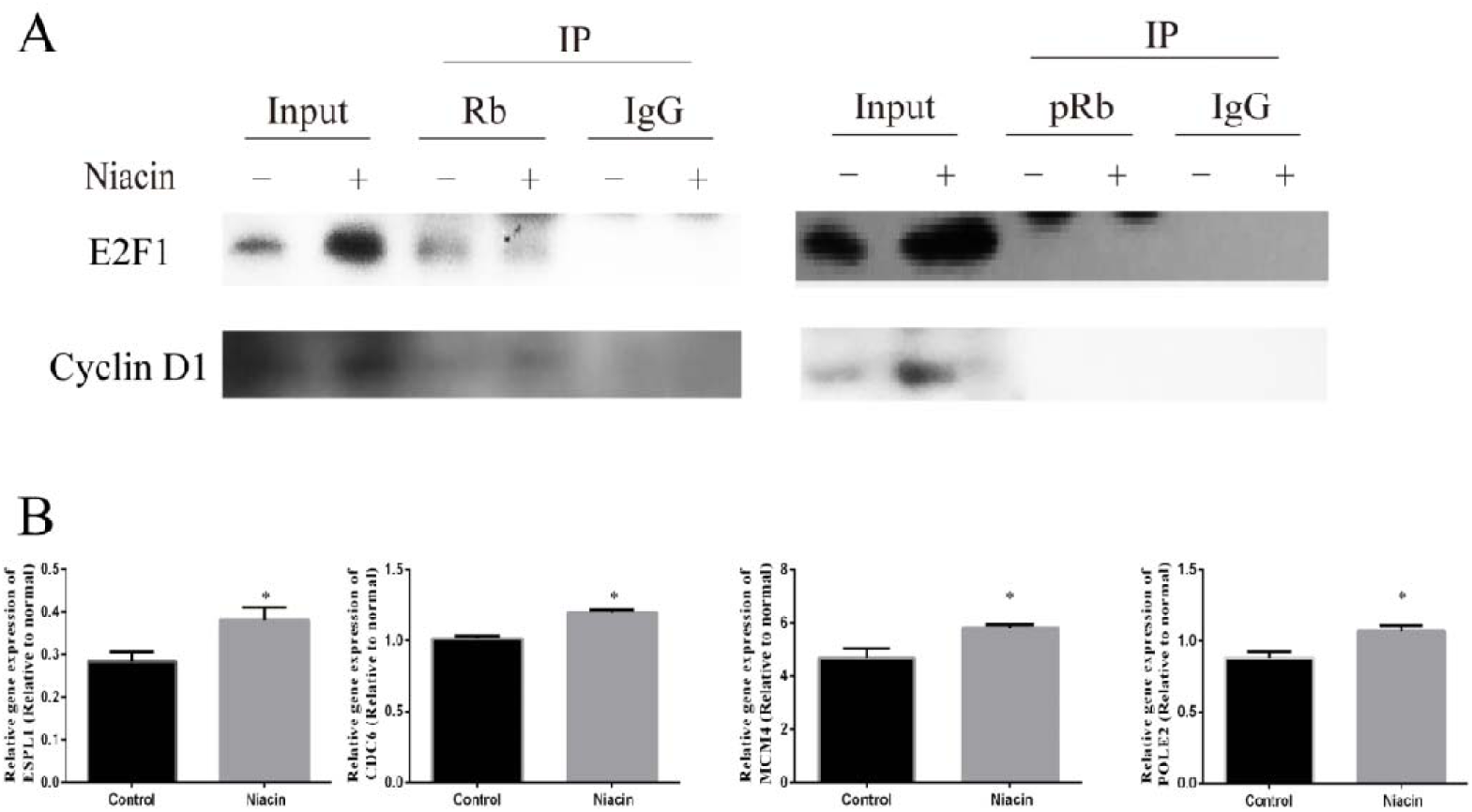
Niacin promotes dissociation of the Rb-E2F1 complex and increases transcription of E2F1 target genes. mMECs were treated with niacin (1 mM) for 2 d. (A) Immunoprecipitation (IP) with anti-pRb, anti-Rb or rabbit IgG (negative control)-conjugated magnetic G beads was carried out in the cell lysates from treated mMECs. Western blots were probed with anti-E2F1 and anti-Cyclin D1. (B) mRNA levels of the E2F1-targeted genes CDC6, ESPL1, MCM4, and POLE2 were detected by real-time quantitative PCR. *P < 0.05 versus the control group.

To confirm that niacin promotes E2F1-targeted genes, a few genes that regulate cell cycle progression and DNA replication were detected by real-time quantitative PCR. The results showed that the mRNA levels of CDC6, ESPL1, MCM4 and POLE2 were significantly increased after niacin stimulation (Figure 7B).

### Peripubertal exposure to drinking water containing 0.5%Niacin promoted mammary duct growth in mice

To test whether niacin promotes mammary gland development in the body, purified water containing 0.5% niacin was fed to 4-week-old weaned mice for 4 weeks. The results showed that the number of ductal branches and terminals increased significantly after drinking water that contained 0.5% niacin (Figure 8A-C). These results suggest that niacin can significantly promote the development of mammary ductal growth in pubertal mice.

**FIG 8.**
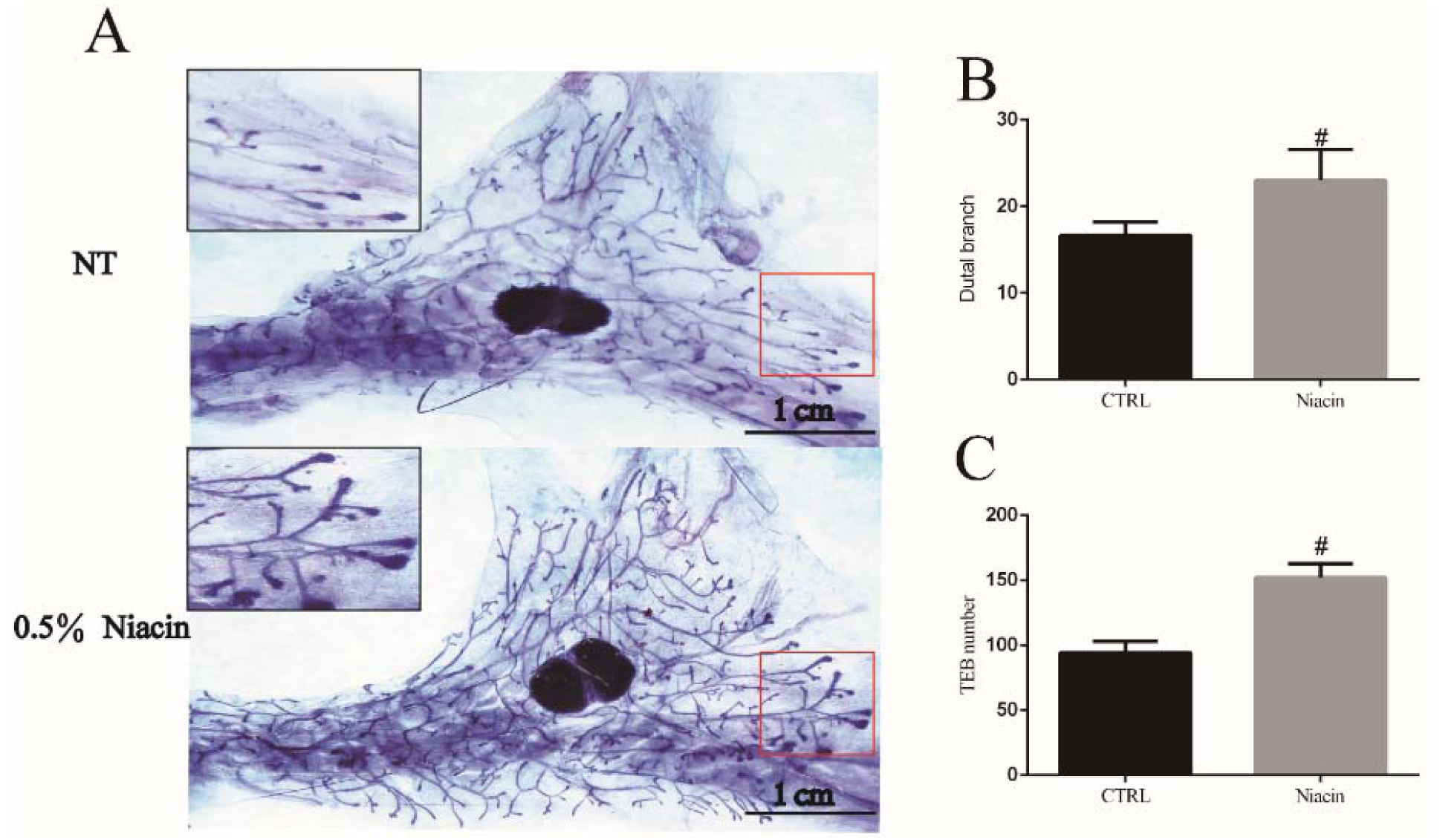
Drinking water containing 0.5% niacin promotes premature mouse mammary duct development. (A) Representative images of whole mount staining of the fourth pair of mammary glands of control and 0.5% niacin-treated pubertal mice. (B, C) Effects of dietary 0.5% niacin on the number of TEBs and ductal branches in the fourth pair of mammary glands of pubertal mice (n = 6). ^#^P < 0.05 versus the control group.

### 0.5% Niacin in Drinking water Promotes Expression of proliferation markers and Activates the AKT/mTOR and ERK Pathways in the Mammary Glands of Pubertal Mice

To further explore the mechanism by which niacin promotes mammary gland development, we found that 0.5% niacin in drinking water significantly increased the phosphorylation levels of AKT, mTOR, ERK and 4EBP1 (Figure 9A, E-H). Moreover, expression of the proliferative marker cyclin D1/D3 and PCNA were also increased by dietary 0.5% niacin (Figure 9A-D). These results suggest that activation of the AKT/mTOR and ERK signaling pathways and subsequent increased expression of cyclin D1/3 and PCNA might be involved in the enhanced mammary gland development induced by dietary 0.5% niacin.

**FIG 9.**
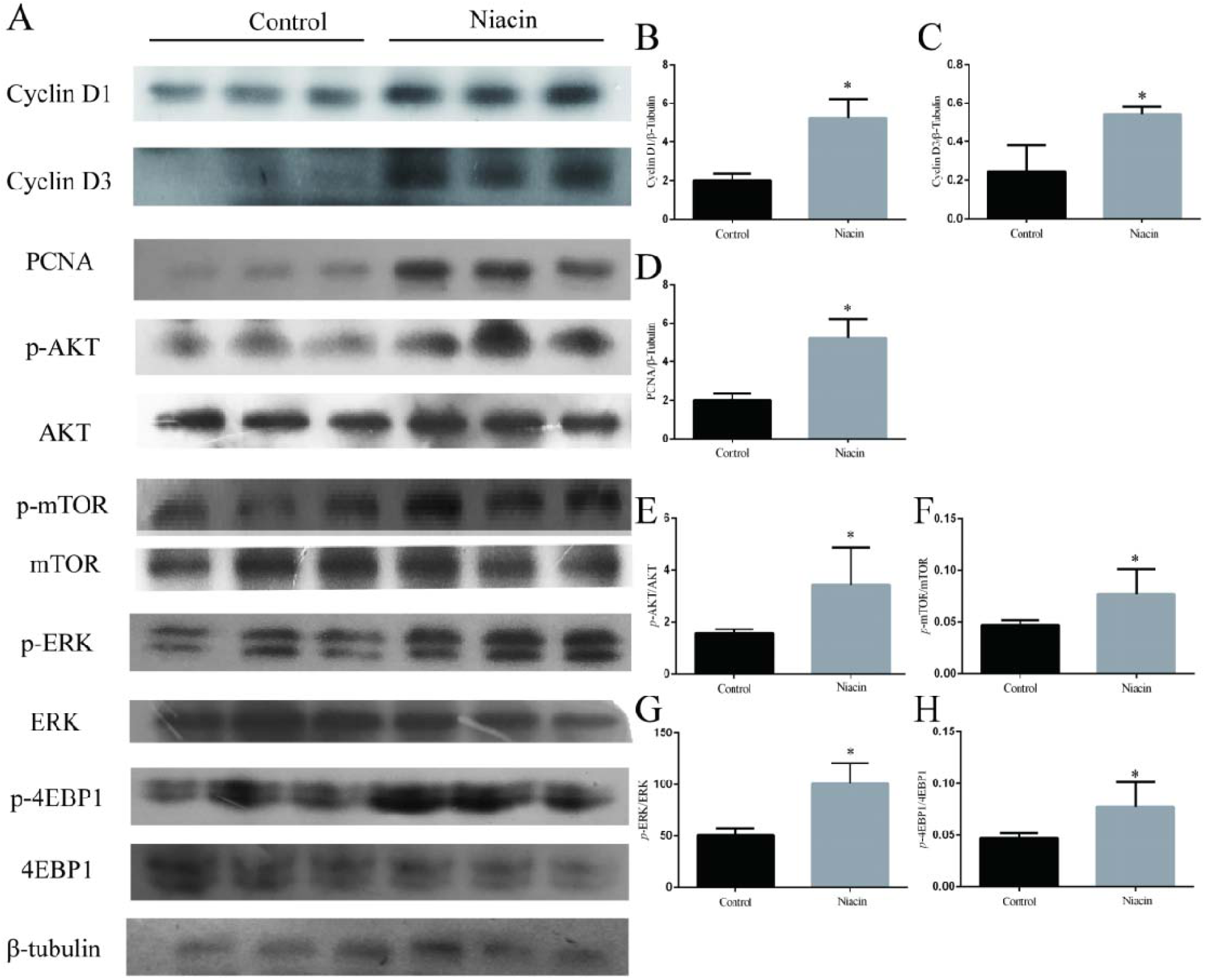
Dietary supplementation with 0.5% niacin increased the expression of cyclin D1, cyclin D3, and PCNA and activated the Akt/mTOR and ERK pathways in the fourth pair of mammary glands of pubertal mice. (A) The protein levels of cyclin D1/D3, PCNA, p-AKT, AKT, p-mTOR, mTOR, p-ERK, ERK, p-4EBP1, and 4EBP1 were examined by Western blotting. (B, C, D, E, F, G, H) Immunoblot bands of Cyclin D1/D3, PCNA, p-AKT/AKT, p-mTOR/mTOR, p-ERK/ERK, and p-4EBP1/4EBP1 were digitized, and cyclin D1/D3 and PCNA were expressed as the ratios to β-tubulin. Data are expressed as the mean ± SEM. *P < 0.05 versus the control group.

## DISCUSSION

In the present study, niacin could activate the AKT/mTOR and ERK signaling pathways by the Gi protein-coupled receptor and increase the phosphorylation level of 4EBP1 to promote the synthesis of cell proliferation markers, leading to dissociation of the Rb-E2F1 complex in mMECs. In addition, 0.5% niacin in drinking water promoted mammary duct development, increased the expression of cyclin D1/D3 and PCNA, and activated Akt/mTOR and ERK1/2 in the mammary glands of pubertal mice.

To our knowledge, this is the first study to assess the effect of niacin on mammary gland development in pubertal mice. It has also been reported that niacin can promote the proliferation of breast cancer cells and cancer stem cells and promote the development of immature oocytes[19-21]. Compared with other vitamins, vitamin A and vitamin D currently inhibit puberty mammary gland development[10, 11]. Niacin promotes the formation of mammary duct branches and terminals in mice. This difference may be due to the different structures of various vitamins. Moreover, we found that niacin also significantly increased the expression of cell proliferation markers (cyclin D1, cyclin D3 and PCNA) in mouse mammary glands. Moreover, niacin also activates the cell proliferation-related signaling pathways AKT/mTOR and ERK1/2. This also suggests that niacin may promote mouse mammary gland development through the AKT/mTOR and ERK1/2 signaling pathways.

Mammary epithelial cells are the most important components of ducts and alveoli, and the central lumen opens to the body surface through the nipple. Most of the epithelial cells are luminal epithelial cells, which are secretory cells that produce milk after functional differentiation during pregnancy[24]. Therefore, the proliferation of mammary epithelial cells is a basic condition for mammary gland development. Cell proliferation markers (cyclin D3, cyclin D1 and PCNA) play an important role in regulating the proliferation of mammary epithelial cells[6-8]. Cyclin D is the core component of the cell cycle mechanism. It activates cyclin-dependent kinases CDK4 and CDK6 and is usually overexpressed in human cancers[25, 26]. Studies have shown that high expression of cyclin D can regulate the proliferation of mammary epithelial cells[6-8]. The results showed that expression of Cyclin D1 and Cyclin D3 in mMECs increased significantly after niacin stimulation. In addition, we found that protein expression of PCNA was also significantly increased. Previous studies have shown that PCNA is an important component of DNA replication machinery, and its expression in S phase is significantly increased, while in stationary and aging cells PCNA expression is low[27]. Subsequent cell cycle results were also consistent with expression of cell proliferation markers. This suggests that niacin may promote the proliferation of mMECs by increasing the expression of cell proliferation markers.

To investigate how niacin promotes proliferation of mMECs through cell proliferation markers, we subsequently examined the proliferation-related signaling pathways AKT/mTOR and ERK1/2. In previous studies, niacin has been shown to protect against brain injury by activating the AKT and ERK1/2 signaling pathways in CHO-K1 and A431 cell lines[16, 28]. In addition, niacin can also inhibit expression of CD40 in LPS-induced HUVECs by activating mTOR[29]. Moreover, activation of the AKT/mTOR and ERK1/2 signaling pathways promoted cell tumorigenesis and proliferation, transformation, migration and invasion of various cancer cells[30, 31]. In this study, niacin significantly activated the AKT/mTOR and ERK1/2 signaling pathways. Subsequent studies with AKT/mTOR and ERK1/2 inhibitors further confirmed that niacin regulates the expression of cell proliferation markers (Cyclin D1/D3 and PCNA) through the AKT/mTOR and ERK signaling pathways. Moreover, inhibition of Gi protein with PTX reversed the niacin-induced activation of the AKT/mTOR and ERK1/2 signaling pathways. These results provided evidence that niacin stimulates mMEC proliferation via the Gi protein-coupled receptor and the linked intracellular AKT/mTOR and ERK1/2 signaling pathways.

To further explore the role of the AKT/mTOR and ERK1/2 signaling pathways in niacin-induced proliferation of mMECs, we used mTOR and ERK inhibitors and detected the activation of 4EBP1. 4EBP1 (also known as EIF4EBP1) is one of the major downstream effectors of translational regulation of mTORC1 kinase. In general, 4EBP1 binds to EIF4E and is immobilized on the 5’ cap end of the mRNA, thereby preventing the formation of translation initiation complexes[32]. Phosphorylation of 4EBP1 by mTORC1 dissociates the 4EBP1-EIF4E complex to form the translation initiation complex. Removing restrictions on protein translation is essential for the development of tissues and many cancers[33, 34]. In this study, the expression level of p-4EBP1 was significantly reduced after stimulation with mTOR or ERK1/2 inhibitors. These results indicate that ERK1/2 promotes 4EBP1 phosphorylation by crosstalk with mTOR in mMECs, thereby promoting the expression of proliferation markers and promoting cell cycle progression. It has been reported that ERK crosstalk with 4EBP1 activates Cyclin D1 translation during quinol-thioether-induced tuberous sclerosis renal cell carcinoma[35, 36]. Therefore, our observations are not special. In addition, we found that niacin also significantly increased the protein expression of p-Rb and E2F1. After adding mTOR and ERK1/2 inhibitors, the expression of pRb and E2F1 decreased significantly, which also indicated that 4EBP1 could regulate the phosphorylation of Rb and the expression of E2F1.

Rb is a key regulator of the G1/S transition[37]. Rb binds to E2F1 to form a complex and inhibits E2F1 activity, thereby inhibiting the transition of cells from G1 to S phase[38]. Cyclin D, which is highly expressed in G1 phase, increases the phosphorylation level of Rb protein by activating CDK and then dissociates the Rb-E2F1 complex[39]. The subsequently dissociated E2F1 can translocate into the nucleus and transactivate its genes to promote cell cycle progression[40]. In this study, niacin-induced expression of cyclin D may be responsible for phosphorylation of Rb protein. Therefore, we used IP experiments to explore the effect of niacin on the Rb-E2F1 complex. The results showed that E2F1 bound to the Rb protein was significantly reduced after niacin stimulation and that cyclin D1 bound to the Rb protein was significantly increased. Moreover, we found that the target gene of E2F1 also increased significantly. These results indicate that niacin promotes proliferation of mMECs by promoting dissociation of the RB-E2F1 complex.

In conclusion, these results indicate that niacin can activate the AKT/mTOR and ERK signaling pathways by Gi protein and increase the phosphorylation level of 4EBP1 to promote the synthesis of celln proliferation markers, which leads to the dissociation of the Rb-E2F1 complex and promotes the development of the mammary duct in puberty mice.

## Author Contributions

Yu Cao and Shoupeng Fu designed experiments. Yu Cao, Lijun Ma, Qing Zhang, Jiaxin Wang, Wenjin Guo, Yanwei Li and Ji Cheng carried out experiments. Yu Cao and Jiafa Wang analyzed sequencing data. Yu Cao, Shoupeng Fu and Juxiong Liu wrote the manuscript.

